# Consistent predictors of microbial community composition across scales in grasslands reveal low context-dependency

**DOI:** 10.1101/2021.11.29.470306

**Authors:** Dajana Radujković, Sara Vicca, Margaretha van Rooyen, Peter Wilfahrt, Leslie Brown, Anke Jentsch, Kurt O. Reinhart, Charlotte Brown, Johan De Gruyter, Gerald Jurasinski, Diana Askarizadeh, Sandor Bartha, Ryan Beck, Theodore Blenkinsopp, James Cahill, Giandiego Campetella, Roberto Canullo, Stefano Chelli, Lucas Enrico, Lauchlan Fraser, Xiying Hao, Hugh A. L. Henry, Maria Hohn, Mohammad Hassan Jouri, Marian Koch, Rachael Lawrence Lodge, Frank Yonghong Li, Janice M. Lord, Patrick Milligan, Hugjiltu Minggagud, Todd Palmer, Birgit Schröder, Gábor Szabó, Tongrui Zhang, Zita Zimmermann, Erik Verbruggen

**Author notes:** **Corresponding author:** Dajana Radujković, University of Antwerp, Universiteitsplein 1, 2610 Wilrijk, Belgium. **Author contributions:** DR and EV conceived the study. DR performed lab work with the help of JDG and DR performed data analysis and interpretation together with EV. DR wrote the first draft of the paper with the help of EV and SV. Other co-authors performed soil sampling and provided information about sites. All co-authors contributed significantly to the final version of the manuscript.

## Abstract

Environmental circumstances shaping soil microbial communities have been studied extensively, but due to disparate study designs it has been difficult to resolve whether a globally consistent set of predictors exists, or context-dependency prevails. Here, we used a network of 18 grassland sites (11 sampled across regional plant productivity gradients) to examine i) if the same abiotic or biotic factors predict both large- and regional-scale patterns in bacterial and fungal community composition, and ii) if microbial community composition differs consistently with regional plant productivity (low vs high) across different sites. We found that there is high congruence between predictors of microbial community composition across spatial scales; bacteria were predominantly associated with soil properties and fungi with plant community composition. Moreover, there was a microbial community signal that clearly distinguished high and low productivity soils that was shared across worldwide distributed grasslands suggesting that microbial assemblages vary predictably depending on grassland productivity.

## Introduction

Variation in the strength and sign of ecological relationships under different environmental, spatial, or ecological settings (i.e. context-dependency) is common in nature (Maestre *et al*. 2005; Chamberlain *et al*. 2014; Tedersoo *et al*. 2015). While biotic and abiotic predictors of microbial community composition have been thoroughly studied at particular spatial scales or environmental contexts (Fierer & Jackson 2006; de Vries *et al*. 2012; Tedersoo *et al*. 2014), it is uncertain whether these predictors are generalizable across different settings. Context-dependency in the processes that structure microbial communities may arise for several (non-mutually exclusive) reasons, including historical legacies (Fukami 2015), stochastic events in community assembly processes (Beck *et al*. 2015), or dispersal limitation (Peay *et al*. 2010), all of which can contribute to the detection of different drivers of microbial community composition depending on region, presence of keystone taxa (Banerjee *et al*. 2018), or environmental conditions (Hendershot *et al*. 2017).

The existence of commonalities in predictors of microbial community composition patterns across sites has been challenging to confirm because most studies have either been restricted in spatial extent or were not designed to evaluate context-dependency. While global-scale studies strongly suggest that a restricted set of predictors such as soil pH (Fierer & Jackson 2006; Delgado-Baquerizo *et al*. 2018) or plant community composition (Prober *et al*. 2015) can universally predict some aspects of soil microbial community composition, the lack of local replication within these global studies complicates distinguishing between different possible drivers that may vary in concert across locations. For instance, microbial and plant communities on the one hand, and soil properties on the other, both strongly covary with geographical distances and climate (Steidinger *et al*. 2019). Regional- and local-scale studies may be better suited to assess the effect of soil properties and plant communities along an environmental (e.g. productivity or fertility) gradient, but findings may not generalize across multiple individual gradients (Alzarhani *et al*. 2019). Indeed, several studies have indicated that the drivers of microbial community composition may strongly vary with spatial and/or environmental contexts (Martiny *et al*. 2011; Shi *et al*.2018; Chalmandrier *et al*. 2019) and that predictability of the soil microbiome depends on spatial scale (Averill *et al*. 2021).

Here, we used a network of 18 grassland sites (containing two to six 64 m^2^ plots; Fig. 1), 11 of which contained plots located along a regional gradient in plant productivity (Fraser *et al*. 2015), to examine the consistency of predictors of soil bacterial and fungal community composition under different spatial scales and environmental contexts. Given that grassland productivity is intrinsically related to biodiversity, soil fertility and plant-soil interactions (Craven *et al*. 2016; Delgado-Baquerizo *et al*. 2017; Guerrero-Ramírez *et al*. 2019), and therefore to the overall ecological functioning of the system, different regional productivity levels provide distinct underlying environmental contexts for the development of soil microbial communities. For instance, plant competition for light is expected to increase with productivity (Grace *et al*. 2016) favouring acquisitive, fast-growing plant species (DeMalach *et al*. 2016) with add-on effects for soils: high input of easily decomposable plant litter selects for more acquisitive microbiota such as many gram-negative and other bacteria (Marschner *et al*. 2011), to the detriment of fungi and microbes engaged in nutritional symbioses with plants (de Vries *et al*. 2007; Johnson *et al*. 2008).

**Figure 1.**
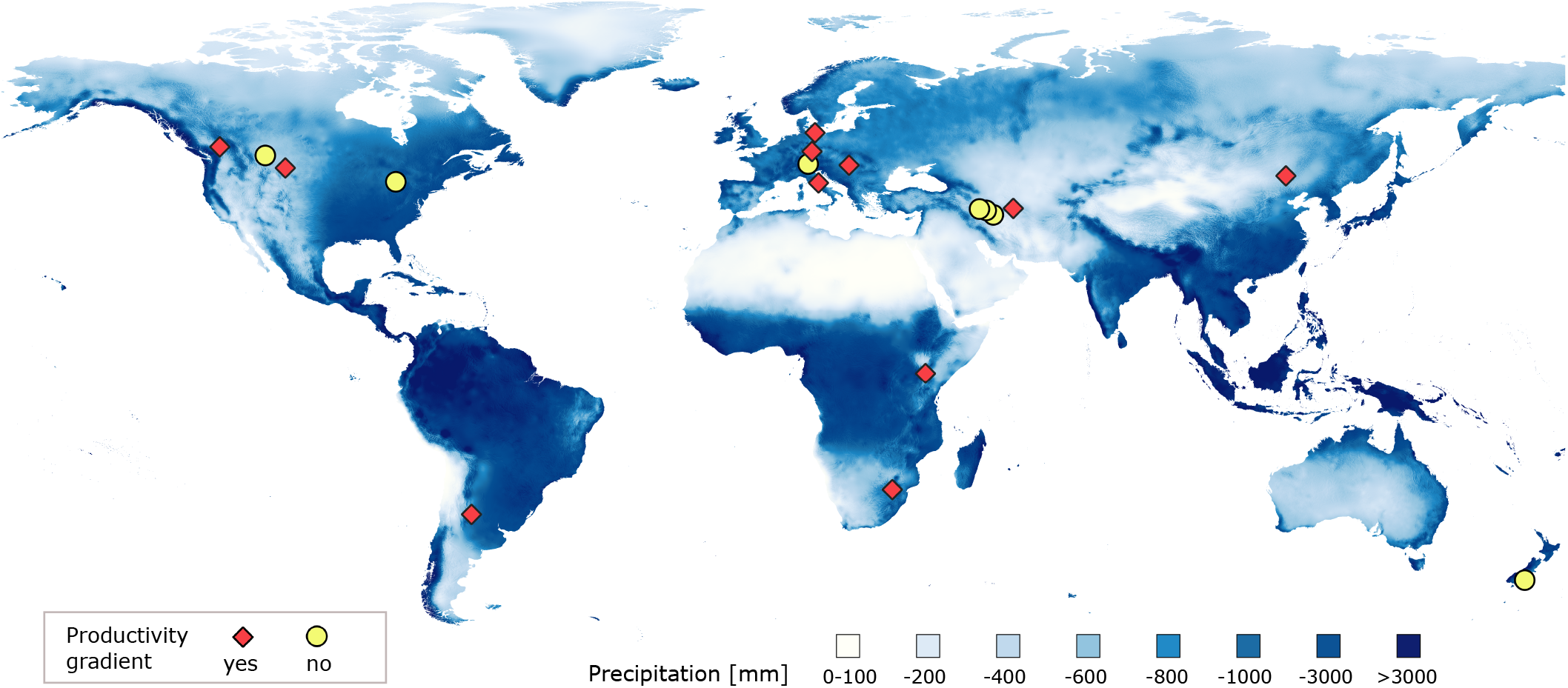
The location of 18 HerbDivNet sites in relation to global precipitation values. Red diamonds indicate 11 sites that contained a clear productivity gradient and yellow circles indicate other sites (containing from 2 to 6 plots but with no clear productivity gradient). All plots (n = 83) were used in the analyses of large-scale predictors of microbial community composition while 11 sites with the productivity gradient (11 pairs of plots with relatively low and high productivity; a total of 44 plots) were used in the analyses of microbial community composition at high and low productivity levels.

To examine whether similar predictors explain variation in microbial community composition across scales, we first analyse the importance of different broad-scale factors (climate, geographical distances, atmospheric nitrogen deposition) and ecosystem fertility-related factors (plant biomass and 14 soil properties) (Table S2) in explaining large-scale bacterial and fungal community dissimilarities. We also test if plant community composition can explain additional variation in microbial community composition when these factors are accounted for. We then examine whether important, regionally-varying, predictors (i.e. ecosystem fertility-related factors and plant community composition) identified at the large scale can likewise consistently predict regional-scale (within-site) microbial community composition, and thus truly ruling out any covariances between sites. Finally, we examine whether two different grassland productivity levels (low and high) have consistent effects on overall microbial community composition across different sites as well as on the correlation networks between major microbial groups, plant functional groups and soil properties. If the drivers of microbial communities are entirely context-dependent, we expect that the important predictors identified at the large scale would be poor or inconsistent predictors of regional-scale variability across sites. Likewise, if the effect of plant productivity on microbial community composition varies strongly across grassland sites (i.e. depending on climatic conditions, biogeography, or soil type), we expect no common signal in microbial community compositional changes between two productivity levels.

## Methods

### Sampling sites and data collection

Data were collected from 18 Herbaceous Diversity Network (HerbDivNet) grassland sites (Fraser *et al*.2015) located in 12 countries distributed over six continents (Fig. 1). The sites include different types of grasslands (xeric, mesic and hydric) spanning a wide range of climatic conditions (mean annual temperature ranges from 1.5 °C to 20.1 °C and mean precipitation ranges from 294 mm to 1237 mm). Peak annual biomass values spanned a range from 13 g/m^2^ to 1187 g/m^2^. Each of the 18 sites contained between two and six plots of 8 × 8 m: 11 sites contained six plots, one site contained four plots, one site three plots and five sites contained two plots (Table S1); to a total of 83 plots. Most sites were chosen to represent a gradient in productivity (low, medium and high; two per each productivity level) with six plots located within the same region with little to no variation in climatic conditions. However, some sites contained fewer plots and did not show a prominent productivity gradient. A clear gradient in biomass productivity was captured in 11 sites; including ten with six plots and one with four plots (Fig. 1).

#### Soil sampling and storage

Soil samples were taken in a single sampling event at the peak of the growing season in the period between 2017 and 2018, depending on the site (Table S1). For each plot within a site, five subsamples were taken from four corners and the centre of the plot at 0-10 cm depth. Subsamples for microbial analyses were taken and stored in pure ethanol (a total of 415 samples) and the rest of the sample was pooled into one composite sample (a total of 83 samples), air-dried and sieved at 2 mm. All samples were further analysed at the University of Antwerp. Samples for microbial analyses stored in ethanol were kept cool until the DNA extraction (see below). Storage in ethanol has been shown to yield similar DNA recovery as cold conservation (Harry *et al*. 2000).

#### Plant sampling

We measured plant species presence and total aboveground biomass from each m^2^ of each 64 m^2^ plot at the peak of the growing season (Table S1). Litter was first excluded from the total biomass and live biomass was dried and weighed. Based on this, average peak biomass production [g/m^2^] was calculated for each plot.

The data on the presence of different plant species at each m^2^ of the plot was used to derive the ‘frequency’ of different species per plot (with the highest possible value of 64 for species present at each m^2^ of the plot) which was used as a measure of relative abundance. Further analyses of plant community composition distances were based on species aggregated to genera (as in Prober et al. (2015)) rather than to the species level because plant species turnover across different plots and sites would often be 100% and thus produce continuous data at highly similar communities only, reducing information content.

#### Climatic, N deposition and soil data

Mean annual precipitation (MAP) and temperature (MAT) were derived from the CHELSA database (Karger *et al*. 2017) based on the geographical position (latitude and longitude) of each plot, which was also used to calculate geographical distances [km] between the plots. Data on total inorganic nitrogen deposition [kg/ha/yr] were derived from Ackerman et al. (2018). We used the average values over the years available in the database to account for long-term fertilization by atmospheric N deposition.

We analysed 14 soil properties: soil organic matter (SOM), total nitrogen (N), total carbon (C), total phosphorus (P), available P, base saturation (BS), cation exchange capacity (CEC), pH, soil texture (sand, clay, silt), extractable Ca, Mg and K. These soil properties are related to soil fertility and plant productivity (Vicca *et al*. 2018), they are known to affect soil microbial community composition (de Vries *et al*. 2012; Tedersoo *et al*. 2014; Zheng *et al*. 2019) and can be compared across different sites. Details on the analyses of soil properties are found in Appendix S1.

### Analyses of microbial communities

#### Sample preparation, sequencing and bioinformatics analyses

DNA was isolated from 415 soil samples using 0.25-0.35 g of soil with the DNeasy PowerSoil Kit according to the manufacturer’s protocol (Qiagen, Venlo, the Netherlands). The bacterial 16S V4 region was amplified using the 515F-806R primer pair (Caporaso *et al*. 2011) and the fungal ITS1 region was amplified using general fungal primers ITS1f (Gardes & Bruns 1993) and ITS2 (White *et al*. 1990), modified according to Smith & Peay (2014). The libraries were sequenced with 2×300 cycles using the Illumina MiSeq platform (Illumina Inc; San Diego, CA, USA). The sequences were analysed using the USEARCH (v8.1.1861) and VSEARCH (Rognes *et al*. 2016) software following the UPARSE pipeline (Edgar 2013) to create operational taxonomic unit (OTU) tables for bacteria and fungi. Representative OTUs were aligned to the SILVA database (bacteria) (Quast *et al*. 2013) (release 138) and UNITE database (fungi) (Kõljalg *et al*. 2005) (release date 2.2.2019), using the *sintax* command in USEARCH with a 0.8 cut-off, resulting in 19,248 and 13,967 OTUs for bacteria and fungi, respectively.

Further steps were performed using R software (R Core Team 2015). The number of reads per subsample was rarefied using the *rrarefy* function in *vegan* (Oksanen & *et al*. 2015) to 6,046 for bacteria and 1,231 reads for fungi as rarefaction curves showed that the number of taxa was levelling off for most subsamples at these depths (Fig. S1). After removing subsamples with too few sequences and/or outliers, there were 402 subsamples for bacteria and fungi (Appendix S1). The sequences from the subsamples were later aggregated to up to five subsamples per plot (see below) so that, overall, plots were represented by up to 30,000 and 6,000 for bacteria and fungi, respectively.

More details on sample preparation, bioinformatics analyses, and fungal functional annotation can be found in Appendix S1.

### Analysis of microbial abundance

DNA extracts of the five subsamples per plot were first pooled into one sample, leaving 83 samples in total. The abundance of bacterial and fungal gene copies per sample was quantified using qPCR targeting 16s V4 region (with the 515F–806R primer pair) for bacteria and 18s region for fungi (primer set FR1 / FF390 (Chemidlin Prévost-Bouré *et al*. 2011)), chosen because high length variation of the ITS1 region precludes accurate quantification. The details on qPCR conditions and quality control are described in Appendix S1.

### Statistical analyses

#### Examining if large-scale predictors consistently explain the regional-scale variation in microbial community composition

We averaged the OTU relative abundances of five subsamples per plot (83 plots in total) to obtain one community measure per plot. Broad-scale (climate, N deposition, geographical distances), ecosystem fertility-related variables (soil variables and plant biomass) and plant community composition were used as potential predictors of large-scale variation in microbial community composition (Table S2). To investigate how well these factors explain the dissimilarities between microbial communities, we created distance matrices using Bray-Curtis (BC) and Euclidean distances, for communities and environmental factors/geographical distances, respectively. All environmental variables (except pH and BS) were transformed using square root transformation, centred and scaled to reduce positive skewness and to allow for the comparison of effect sizes. Community data (fungi, bacteria, plants) were transformed with Hellinger transformation using the *decostand* function in the *vegan* package in R.

The influence of different factors on the dissimilarity in bacterial/fungal communities was analysed using multiple regression on distance matrices (MRM) in the *ecodist* package (Goslee & Urban 2007). MRM model was first fitted using bacterial/fungal distances as response variables and broad-scale and ecosystem fertility-related environmental variables as predictors. The variables that did not significantly contribute to the model were removed leaving only the variables with a significant effect (P < 0.05). This was done to comprehensively capture the effect of the environment (and geographical distances) on microbial community composition and to retrieve the effect sizes of different important variables that were later used to construct regional-scale environmental variable (see below). To test if plant community distances can explain any unique (non-shared) variation in microbial community composition, we included it in the model with broad-scale and ecosystem fertility-related variables and we partitioned the variation explained by these three groups of variables. Therefore, given that microbial and plant community distances can be related due to shared environmental conditions, we accounted for a vast number of environmental variables (without necessarily attempting to disentangle the effect of different correlated environmental predictors) before assessing if plant community composition explains additional variation in microbial community composition.

To examine if the observed large-scale relationships (across all the plots and all the sites) persist at the regional scale (i.e. between the plots within each site, which share a similar climate and are part of the same species pool), we created a common variable that represents the influence of the important ecosystem fertility-related variables by first multiplying each variable by its coefficient in the MRM large-scale model and then summing them. In this way, we were able to ‘weigh’ the importance of different fertility-related variables (while accounting for climate and geographical distances) and test if the resulting ‘environmental variable’ can consistently explain the regional-scale variation in microbial community composition. The within-site (Euclidean) distances in the environmental variable were then regressed against the within-site distances in bacterial and fungal communities. Finally, the within-site microbial distances were also regressed against the within-site plant community distances to examine how well plant community dissimilarities can predict microbial community dissimilarities at the regional scale. To assess the consistency of these relationships (environment – bacteria, plants – bacteria, environment – fungi, plants – fungi) across sites that contained more than three plots, we calculated the variance in their slope values and reported their mean R^2^ values and standard deviations.

#### Microbial community composition at different regional relative productivity levels

Our regional productivity gradients allowed us to test whether there is a general difference between relatively low-productivity and high-productivity grasslands replicated at large scale. For this analysis, the dataset was divided into two subsets: one containing two plots with low productivity and the other containing two plots with high productivity from each site. Eleven sites with a clear productivity gradient were selected yielding two datasets each containing 22 plots. These sites had a strong difference in plant biomass between the plots of low and high productivity (two plots with high productivity within a site had on average at least 100% higher biomass than those with low productivity).

To test if bacterial and fungal communities differed significantly between the two productivity levels with a consistent pattern across globally distributed sites, we performed PERMANOVA analysis using the *adonis* function in *vegan* adding ‘site’ as *strata* to control for inherent community differences between sites. We used multidimensional scaling (MDS) ordination to visualise the BC distance in bacterial and fungal communities at different productivity levels after removing the effect of ‘site’ differences using the *dbrda* function in *vegan*. To examine if the best predictors of bacterial and fungal community composition differed at different productivity levels, we repeated the model selection described above (using the MRM function) for microbial communities for each of the productivity levels. Furthermore, using the *multipatt* function (with 999 permutations) from the *indicspecies* package, we determined bacterial and fungal OTUs which were significant (P < 0.01) indicators of low and high productivity levels. We also examined if there was a significant difference (P < 0.01) in the relative abundances of bacterial and fungal groups (taxonomic and functional, respectively) and total bacterial and fungal abundances (number of gene copies) at low compared to high productivity levels using the *lme* function in *nlme* package with ‘site’ as a random effect. The normality of residuals was tested using the Shapiro-Wilk test.

Finally, we examined whether the correlation networks between microbial groups/total microbial abundances, plant functional groups and soil properties across different sites differed between low and high productivity levels. To this end, we analysed the pairwise correlations (using *corr.test* in the *‘psych’* package) between the three most dominant bacterial phyla, three most dominant fungal functional groups, three plant functional groups (grasses, forbs, legumes), fungal and bacterial abundances, plant biomass and the most important soil properties (SOM, CEC, BS, pH, total N, C:N, total P, available P and % sand), for low and high productivity datasets. Only the correlations with Spearman r > 0.5 and P-value < 0.01 were retained and visualised in the form of correlation networks.

## Results

### Predictors of microbial community composition at large vs regional scale

Our results revealed that a composite environmental variable created using the most important fertility-related variables in the large-scale model (with the strongest effect of base saturation and pH; Table S3) consistently predicted regional-scale (within-site) variation in bacterial community composition across sites (slope variance = 0.05; mean R^2^ = 0.58, sd = 0.32) (Fig. 3a). Plant community composition explained additional variance in bacterial community composition at the large scale after important broad-scale and ecosystem fertility-related variables were accounted for (explaining more unique variation than broad-scale predictors, Fig. 2). At the regional scale, plant community composition was also consistently and strongly associated with the variation in bacterial community composition for most sites (slope variance = 0.06; mean R^2^ = 0.64, sd = 0.28) (Fig. 3b).

**Figure 2.**
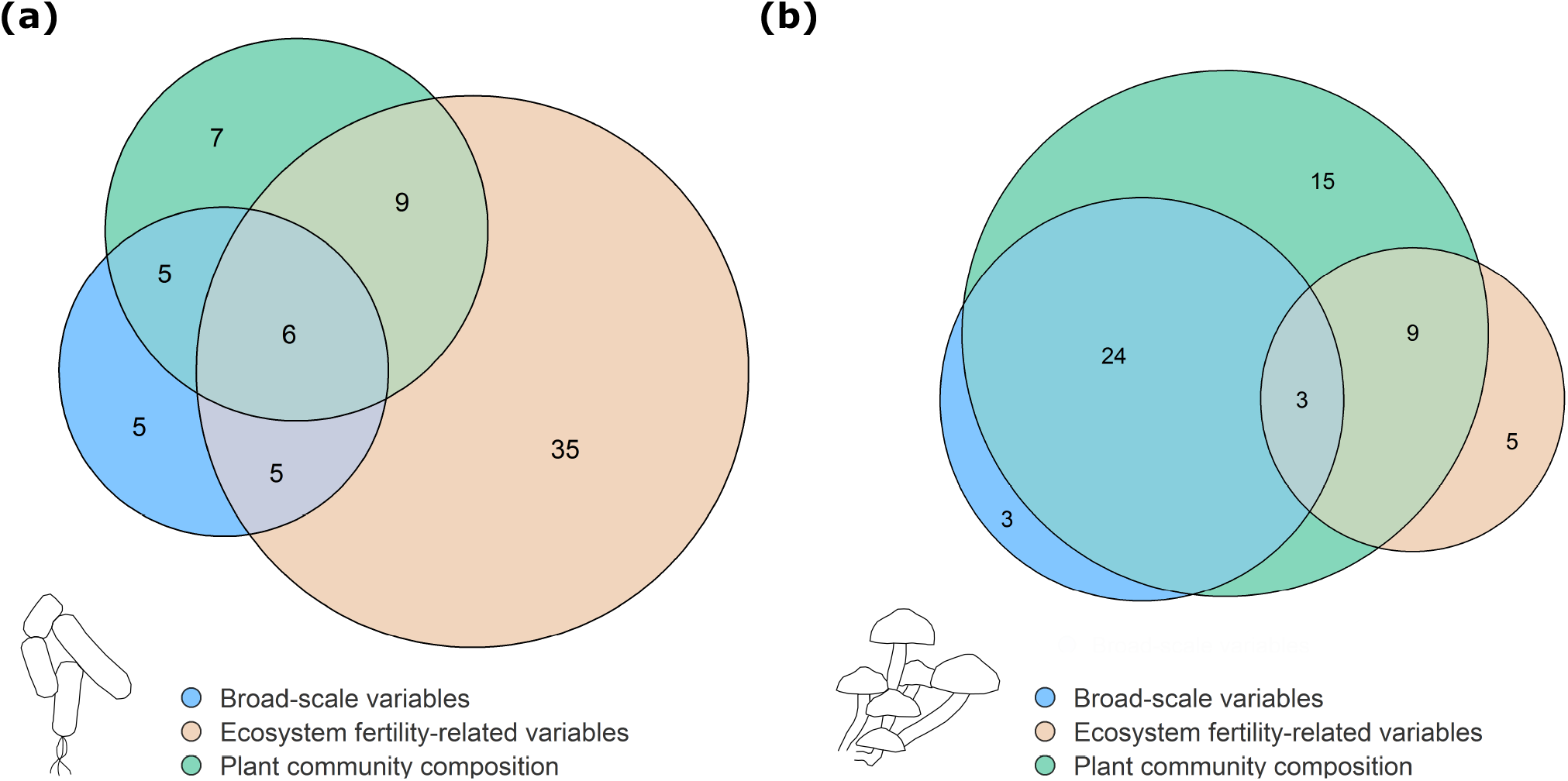
Variance partitioning between selected variables in the large-scale model explaining a) bacterial and b) fungal community composition. The variables were grouped in three categories: i) broad-scale variables (climate, N deposition and geographical distance); ii) ecosystem fertility-related variables (soil properties and biomass) and iii) plant-community composition. The sizes of bubbles correspond to the percentage of variance explained by each group (indicated by the numbers in the bubbles).

**Figure 3.**
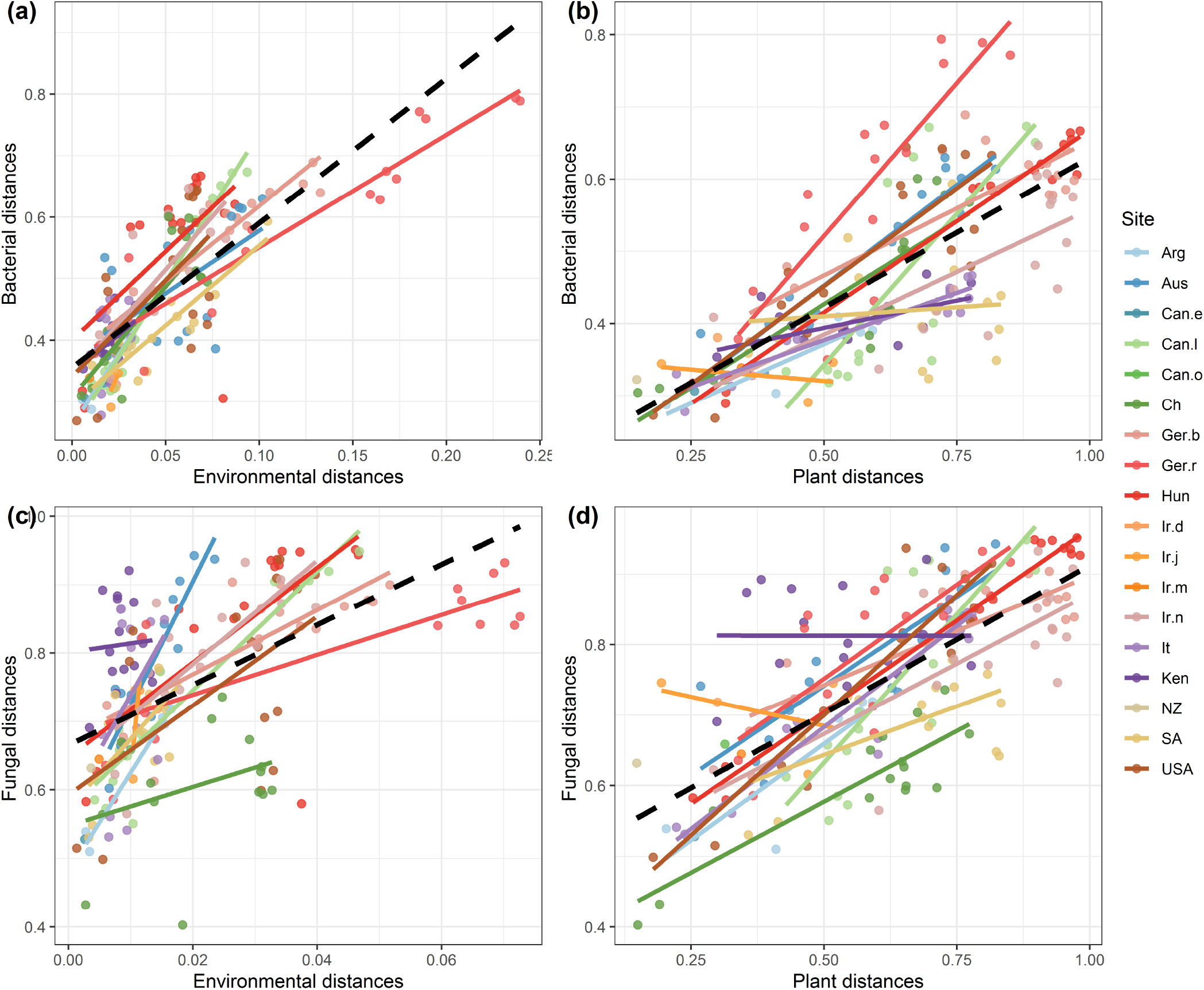
Relationships between regional (within-site) environmental/plant community distances and bacterial and fungal community distances **a)** bacterial distances vs environmental distances; **b**) bacterial distances vs plant distances **c**) fungal distances vs environmental distances; **d**) fungal distances vs plant distances. Colours of points and corresponding regression lines correspond to 18 different sites. Dashed lines represent general regression lines. The relationship between regional geographical distances and bacterial/fungal distances per site are shown in Fig. S2. For site references, see Table S1.

The consistency between large- and regional-scale predictors was found for fungi as well, where the best large-scale predictor (plant community composition) was also consistently associated with the within-site variation in fungal community composition for most sites (slope variance = 0.05; mean R^2^ = 0.64, sd = 0.26) (Fig. 3d). Plant community composting was a better predictor at the large-scale than all broad-scale and ecosystem fertility-related variables combined (R^2^ = 0.51 and R^2^ = 0.44, respectively) (Table S3, Fig. 2). Accordingly, the relationship between fungal community composition and the composite environmental variable varied considerably from site to site (slope variance = 0.16; mean R^2^ = 0.50, sd = 0.32) (Fig. 3c).

### Microbial community composition at different plant productivity levels

Bacterial and fungal community composition differed significantly (P < 0.001) between the two productivity levels (Fig. 4) when site differences were accounted for. This indicates that there is a common community, shared across the globally distributed sites, which can separate more and less productive grasslands. Despite the compositional differences, the predictors of microbial community composition at low and high productivity levels were similar. In line with the results in the previous section, soil properties (particularly base saturation and pH) were the most important predictors of bacterial community composition, whereas fungal community composition was best predicted by plant community composition (Appendix S2). Therefore, while distinct microbial communities were found at contrasting productivity levels, their associations with the abiotic or biotic environments across sites were largely similar.

**Figure 4.**
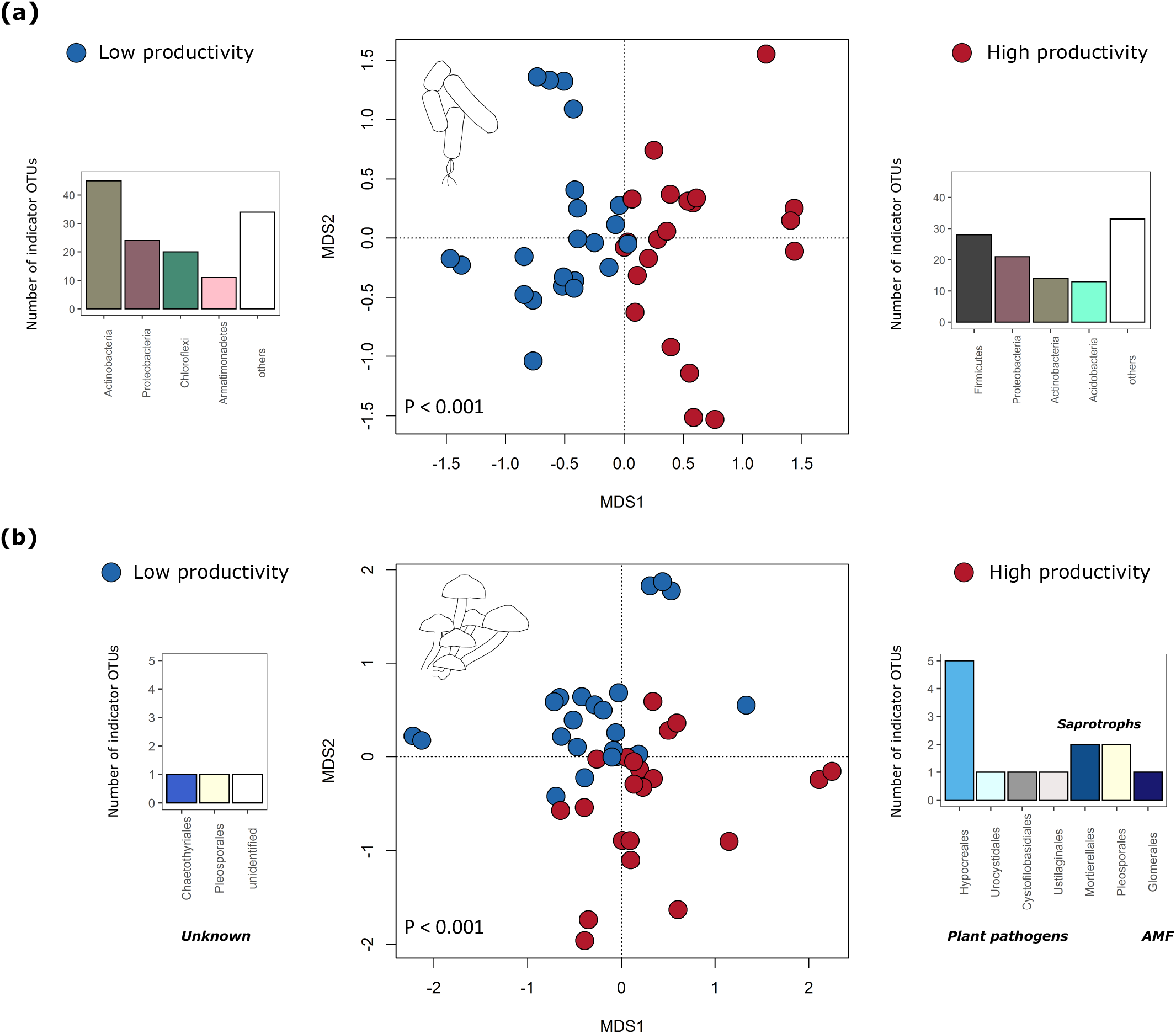
Partial MDS ordination showing **a)** bacterial and **b)** fungal Bray-Curtis distances (partialling out the effect of the site differences) coloured according to the productivity level of the sampling plots. The bar plots on the sides present **a)** the number of bacterial OTUs split by phylum and **b)** fungal OTUs split by order, which were found to be significant indicators of low and high productivity grassland soils. For fungi, putative trophic lifestyles of these OTUs are indicated in bold.

To further disentangle the effect of different productivity levels on microbial communities, we examined the most important bacterial phyla and fungal functional groups. The most abundant (> 10% relative abundance) bacterial phyla in the dataset were: Actinobacteria (42%), Firmicutes (16%) and Proteobacteria (14%) (Fig. S3, Table S4). Saprotrophs were the most dominant fungal functional group with 54% of sequences followed by 14% of potential plant pathogens, 7% of arbuscular mycorrhizal fungi (AMF), whereas the other groups together accounted for 4% of the total number of sequences (Fig. S3).

We further used indicator species analysis to identify the OTUs that significantly associate with different productivity levels. There were 109 and 134 bacterial OTUs indicators of high and low productivity sites, respectively. The highest number of indicators for low productivity belonged to Actinobacteria (33.6%; dominant order was Thermoleophilia) while for high productivity, they predominantly belonged to Firmicutes (25.7%), many of which were from the order Clostridia (22.9%) (Fig. 4a). In the case of fungi, the high productivity sites had 13 indicators, most of which were assigned as putative plant pathogens, predominantly from the Nectariacea family (smut fungi). On the other hand, low productivity sites had only 3 indicator OTUs whose trophic lifestyle was unassignable at the genus level (Fig. 4b).

When considering total bacterial and fungal abundance (number of gene copies) and the three most dominant fungal and bacterial groups, the linear mixed-effect model with ‘site’ as a random effect showed that Actinobacteria and total fungi were more abundant in low than in high productivity sites, and the opposite was observed for Firmicutes (Fig. 5a). The relative abundances of Proteobacteria, saprotrophs, AMF and total abundance of bacteria, did not differ significantly between the two productivity levels.

**Figure 5.**
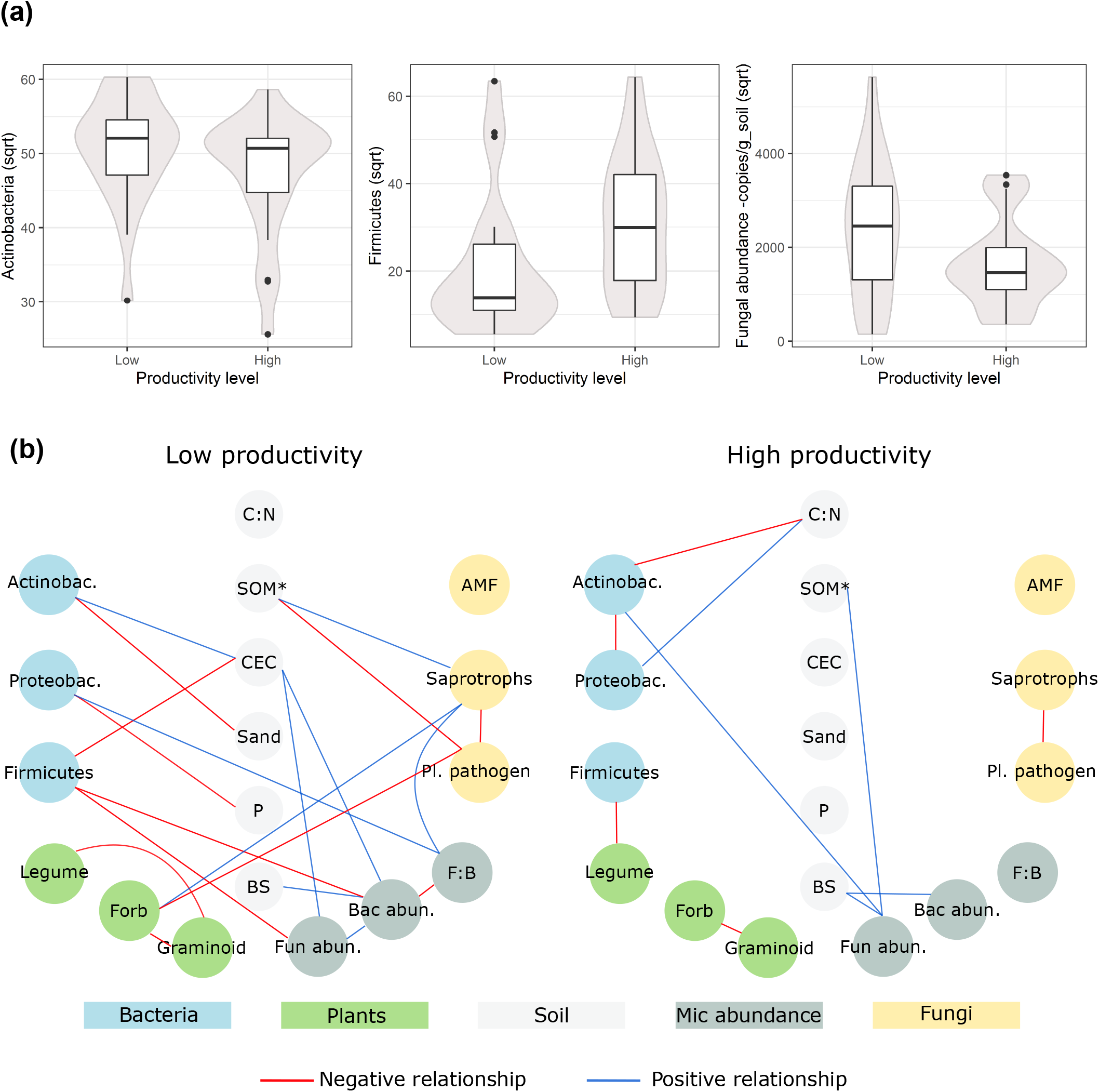
**a**) Boxplots showing the mean values of Actinobacteria and Firmicutes relative abundances and total fungal abundances at two productivity levels (the differences are significant in all cases). The grey area depicts the distribution of samples. **b**) Correlation networks between the three most dominant bacterial phyla (Actinobacteria, Firmicutes, Proteobacteria), three dominant fungal functional groups (saprotrophs, putative plant pathogens, AMF), three main plant functional groups (grasses, forbs, legumes), total bacterial/fungal abundance (number of copies per g soil) and soil properties at high and low productivity. Soil variables that had at least one significant correlation are shown. The red lines depict significant negative correlations, while blue lines depict significant positive correlations (P < 0.01 and Spearman r > 0.5). Soil variables included C:N (carbon to nitrogen ratio), N (total nitrogen), CEC (cation exchange capacity), percentage sand, P (available phosphorus), BS (base saturation). SOM* = the same links were observed for total N and P, which were all strongly correlated to each other and only one of them is shown. The correlations between soil properties are not shown.

Although microbial community composition was explained by similar predictors at low and high productivity grasslands, at higher levels of taxonomic and/or functional integration this was not the case. The correlation networks between the three most dominant bacterial and fungal groups with plant functional groups (graminoids, herbs and legumes), soil properties and total fungal and bacterial abundance differed substantially between the two productivity levels. At high productivity, there were only a few correlations; e.g. between C:N and both Actinobacteria and Proteobacteria. On the other hand, the number of associations was much higher at the low productivity level (Fig. 5b) where different soil properties were associated with fungal and bacterial groups. Moreover, there were negative correlations between putative plant pathogens and forbs as well as between Firmicutes and total bacterial and fungal abundances.

## Discussion

Despite considerable literature describing the most important predictors of soil microbial community composition in the grassland biome, until now it has been unclear whether these relationships persist across different spatial and environmental contexts. In this study, we show that there is generality in the way bacterial and fungal communities are shaped across two different spatial scales and productivity levels in worldwide distributed grasslands.

### Generality in the predictors of microbial community composition

Our results reveal that soil abiotic factors (primarily base saturation and pH) are key predictors of bacterial community composition both across and within different grassland sites and at contrasting plant productivity levels. The potential role of soil chemical properties (i.e. soil pH) as important drivers of continental-scale bacterial community turnover (Fierer & Jackson 2006; Lauber *et al*. 2009), as well as of globally dominant bacterial phylotypes (Delgado-Baquerizo *et al*. 2018) has previously been established. However, besides soil properties, bacterial community composition was also strongly and consistently associated with plant community composition, particularly at the regional scale. These results suggest that at the regional scale, plant community composition and soil chemical properties might jointly influence bacterial communities and their individual importance may be difficult to disentangle. Fungal community composition was consistently related only to plant community composition, indicating that plant communities, rather than soil properties (Egidi *et al*. 2019), are important in shaping fungal community composition in grasslands.

Large-scale association between grassland plant community composition and both fungal and bacterial community composition has previously been demonstrated (Prober *et al*. 2015). The consistency of the relationship between plant and microbial (particularly fungal, but also bacterial) community composition across different grasslands in our study shows that these relationships are not just a matter of coincident spatial community turnover between fungi (bacteria) and plants, but rather indicate a direct influence on each other and/or a high similarity in ecological niches. Plant communities can affect soil microorganisms both directly by providing a diverse set of hosts for mutualistic and antagonistic microorganisms, and indirectly by altering edaphic factors and providing different quantity and quality of root exudates and litter (Wardle *et al*. 2004; Van Der Heijden *et al*. 2008; Berg & Smalla 2009). Local experiments have previously shown that plant community composition can shape microbial community composition (Schlatter *et al*. 2015; Reese *et al*. 2018; Heinen *et al*. 2020) and that plant-microbe feedbacks might play a central role both in microbial and plant community assembly processes (Wubs *et al*. 2019; Radujković *et al*. 2020).

### Universal influence of plant productivity on soil microbial community composition

Bacterial and fungal community composition were found to be more similar within low and high productivity grasslands than between them when site-specific differences were accounted for. This suggests that plant productivity as an indicator of a myriad of factors related to it (including soil fertility, plant diversity, and plant-soil interactions (Craven *et al*. 2016; Delgado-Baquerizo *et al*. 2017; Guerrero-Ramírez *et al*. 2019)) selects for some of the same microbial taxa regardless of differences in climate and grassland type. A link between bacterial taxa and plant productivity across contrasting biomes worldwide (forests, shrublands, grasslands) has previously been reported (Delgado-Baquerizo *et al*. 2018), where particular groups of globally dominant soil bacteria with a preference for low-productive sites were identified. Here, we show that similar conclusions hold for bacterial and fungal taxa even within the grassland biome, where differences in plant productivity are much smaller than across contrasting biomes.

The differences in bacterial community composition between the two productivity levels in our study are corroborated by a higher relative abundance of Firmicutes and lower relative abundance of Actinobacteria at high productivity. OTUs belonging to the phylum Firmicutes were also found to be the most dominant indicators of high productivity soils. This is consistent with the findings of several other studies showing an increase in Firmicutes abundance under elevated nutrient inputs suggesting that many members of this phylum may be associated with fertile soils (Ramirez *et al*. 2010; Wakelin *et al*. 2013; Yao *et al*. 2014; Ling *et al*. 2017). Among the indicators of low-productivity grasslands, many belonged to the phylum Actinobacteria, particularly the order Thermoleophilia. Members of this order are known to thrive in conditions of reduced soil moisture (Pereira de Castro *et al*. 2016; Ochoa-Hueso *et al*. 2018; Preece *et al*. 2019) which might explain their presence in low-productivity grasslands with their predominantly sandy soils and poor water-holding capacity.

The relative abundances of the three dominant fungal functional groups (saprotrophs, AMF and putative plant-pathogens) did not differ significantly between productivity levels. However, total fungal abundance was significantly higher at low compared to high productivity levels. Higher fungal abundance is common in less fertile soils (Bardgett & McAlister 1999; Innes *et al*. 2004) where fungi are favoured over bacteria as the predominant decomposers due to the higher recalcitrance of plant litter and their generally more resource-conservative lifestyles (Marschner *et al*. 2011). Moreover, plant reliance upon, and allocation to AMF is often higher to secure P, N and other nutrients (Verbruggen *et al*. 2013; Ven *et al*. 2019). Most of the indicators of highly productive grassland soils belonged to the groups of putative plant pathogens. Plant pathogens are known to thrive under the conditions of high productivity (Reynolds *et al*. 2003) and our result suggests that some of their members are broad generalist appearing in different highly productive grasslands. Low-productivity grasslands appear to share few fungal taxa, possible because these grasslands are more heterogeneous with higher levels of endemism.

### The associations between microbial groups and the environment vary with plant productivity

We explored the factors that potentially drive the total microbial abundances and relative abundances of dominant, bacterial taxonomic groups and fungal functional groups at different productivity levels. Microbial groups from low-productive soils were significantly correlated with many more environmental factors (either plant functional groups or soil properties) than those from high-productive soils. For instance, at low productivity, the relative abundance of putative plant pathogens was negatively associated with the abundance of forbs and tended to increase with increasing graminoid abundance. The tendency of graminoids to accumulate fungal pathogens relative to forbs is a commonly observed phenomenon (Heinen *et al*. 2020) and may be related to their typical high density (Mitchell *et al*. 2002). At the high productivity level plots, plant pathogens and saprotrophs were not correlated with other groups of biota or with soil properties, possibly indicating relaxation of biotic/abiotic interactions when resources are abundant.

These examples suggest that microbial groups at high productivity plots might not be substantially affected by a further increase in resource availability and they might be forming fewer consistent interactions (symbiotic or competitive) with each other or with plant groups. This has been demonstrated in agricultural settings where fertilization reduced rhizosphere microbiome dependency on plant-derived carbon leading to simpler plant-microbe associations (Ai *et al*. 2015). Similarly, it has been shown that 150 years of fertilization has weakened the complexity of plant-microbiome networks in a managed grassland (Huang *et al*. 2019). Our results support that these tendencies also appear to hold for non-agricultural grasslands. Therefore, bacterial taxonomic and fungal functional groups (and by extension, the functions performed by these groups) in low-productivity grasslands may be more strongly influenced by changes in soil properties and plant functional groups than those in high-productivity grasslands.

## Conclusion

Universal ecological patterns are the exception rather than the rule (Lawton 1999) and several studies have argued that there are few if any, general drivers of microbial community composition. If estimates derived from one system or spatial scale cannot be extrapolated to another, it is challenging to predict the effects of altered environmental conditions on soil microbial communities and the functions they drive. Our findings suggest that the main factors that shape overall microbial (bacterial and fungal) community composition in grasslands agree in a highly consistent manner, regardless of the spatial scale, productivity, or climatic conditions while the drivers of the (relative) abundance of specific bacterial and fungal groups may depend on grassland productivity. Moreover, particular, regional productivity levels are typified by relatively similar soil microbial communities across the grassland biome and are distinguishable by that characteristic. These findings suggest that it is possible to extrapolate and upscale the general trends regarding the drivers of microbial community composition and that modelling soil microbial community composition under environmental changes, or using microbial fingerprints to distinguish fertile from infertile systems, are feasible tasks.

## Acknowledgements

This research was supported by the Research Foundation—Flanders (FWO), the European Research Council grant ERC-SyG-610028 IMBALANCE-P and Methusalem funding of the Research Council UA. BS, ZZ and GS were supported by the GINOP-2.3.2-15-2016-00019 project. We thank Annelies Devos, Tom Van der Spiet, Kevin Van Sundert and Irene Ramirez Rojas for their help with the laboratory analyses of soil samples. GJ wants to particularly mention the contribution of Marian Koch, who died suddenly and unexpectedly in April 2020. Marian was a wonderful person and a dedicated vegetation ecologist who organized and lead the fieldwork at the site Ger.r. We miss him.

