## Supplementary Information for "Consistent predictors of microbial community composition across scales in grasslands reveal low context-dependency"

Supporting information

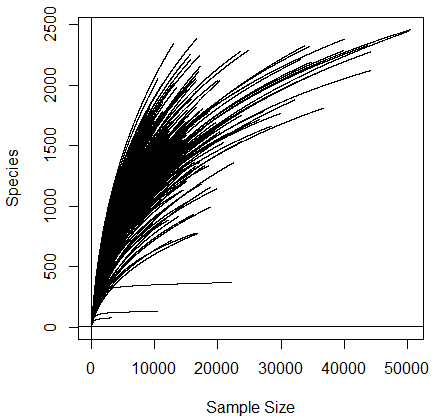

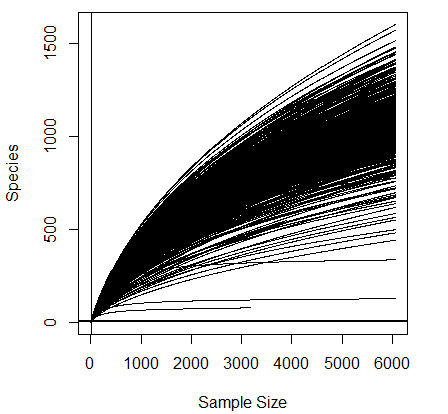

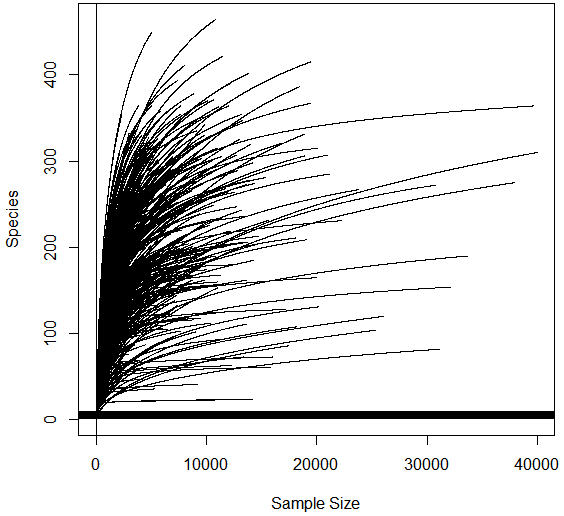

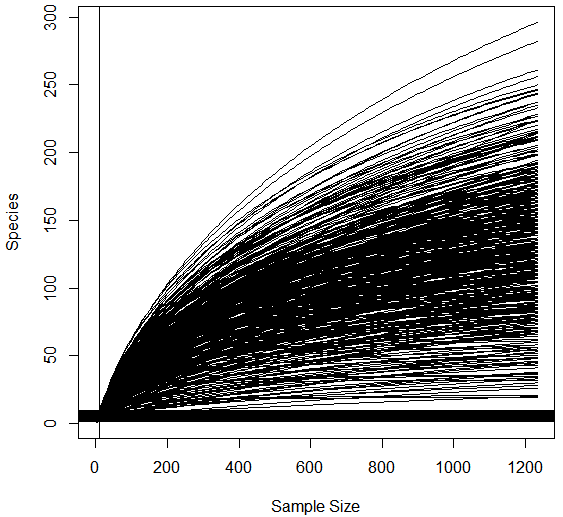

**Figure S1** Rarefaction curves for bacteria (top) and fungi (bottom). The graphs on the left show the curves for non-rarefied data, while the graphs on the right show the curves after rarefactions at 6046 and 1231 reads for bacteria and fungi, respectively.

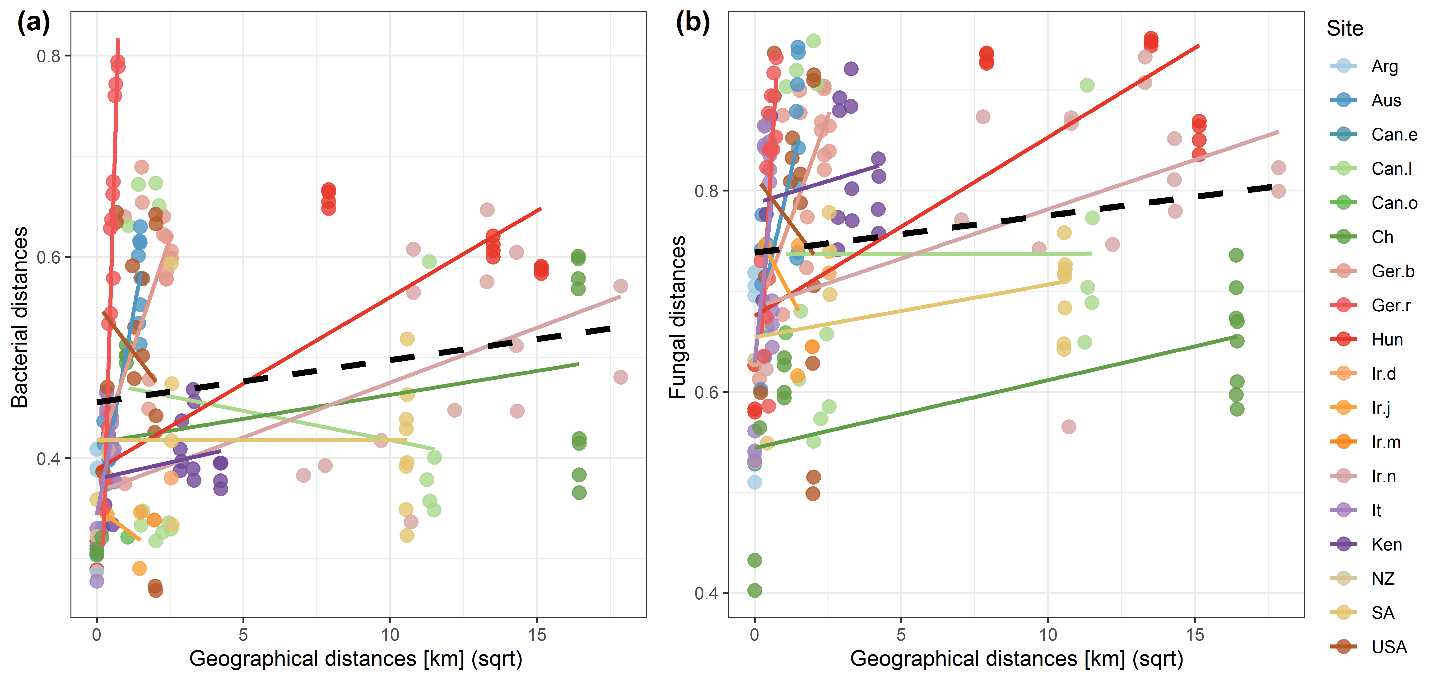

**Figure S2** The relationships between within-site geographical distances and within-site BC distances in a) bacterial and b) fungal community composition. Out of the six sites that have relatively high distances between some plots (between 10 and 300 km), the relationship was found to be relatively strong (R^2^ ranging from 0.3 to 0.57) for only one site in the case of bacteria (Hun) and two sites for fungi (Hun and Ch). In all cases, plant community composition explained a much higher proportion of the variation in microbial communities (R^2^ between 0.77 and 0.98). This demonstrates that within-site geographical distance is not the main cause for the observed relationships between environmental variables/plant community composition and microbial community composition shown in Fig. 3. Site references as in Table 1.

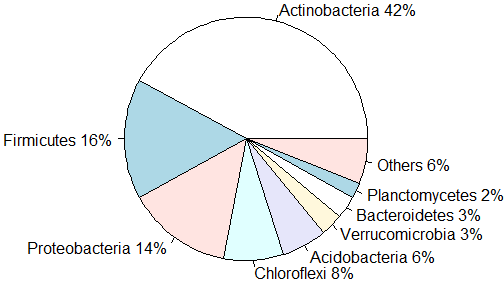

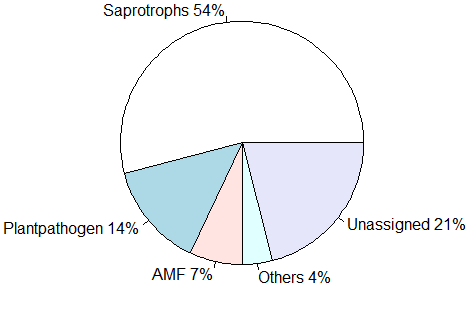

**Figure S3** The percentage of total reads belonging to particular bacterial phyla (top) and fungal functional groups (bottom) in all the samples included in the study.

**Table S1** Information about 18 sites included in the analyses. The sites shaded in grey contained a productivity gradient (i.e. they contained 2 pairs of plots located in the same region and exposed to similar climatic conditions but with a more than 2-fold difference in biomass production).

| ***Site ID*** | ***Site name*** | ***Country*** | ***# of plots*** | ***Year: biomass sampling*** | ***Year: soil sampling*** | ***Mean MAP (sd)*** | ***Mean***  ***MAT (sd)*** |
| --- | --- | --- | --- | --- | --- | --- | --- |
| Aus | Pitztal | Austria | 6 | 2012 | 2017 | 796 (50) | 4.6 (0.9) |
| Arg | El Cuadrado, Córdoba | Argentina | 4 | 2012 | 2017 | 995 (0) | 12.9 (0) |
| Can.e | Elginfield Observatory | Canada | 2 | 2012 | 2018 | 1001 (0) | 7.9 (0) |
| Can.l | Lac du Bois | Canada | 6 | 2012 | 2018 | 379 (18) | 6.5 (0.5) |
| Can.o | Onefour, AB | Canada | 2 | 2012 | 2017 | 301 (10) | 6.7 (0.1) |
| Ch | Inner Mongolia | China | 6 | 2017 | 2018 | 207 (82) | 2.1 (0.6) |
| Ger.b | Bayreuth | Germany | 6 | 2012 | 2017 | 721 (38) | 8.5 (0.2) |
| Ger.r | Rostock | Germany | 6 | 2015 | 2017 | 668 (0) | 8.7 (0) |
| Hun | Soroksár, Fülöpháza, Battonya | Hungary | 6 | 2012 | 2018 | 561 (5) | 11 (0.4) |
| Ir.d | Damavand | Iran | 2 | 2012 | 2017 | 425 (16) | 7.0 (0.5) |
| Ir.j | Javaherdeh | Iran | 3 | 2012 | 2017 | 1230 (13) | 5.3 (1.3) |
| Ir.m | Masuleh | Iran | 2 | 2012 | 2017 | 843 (54) | 5.5 (0.1) |
| Ir.n | North Khorasan, Golestan | Iran | 6 | 2012/2017 | 2017 | 398 (65) | 8.6 (2.5) |
| It | Torricchio Nature Reserve | Italy | 6 | 2012 | 2017 | 1057 (21) | 8.3 (0.1) |
| Ken | Laikipia | Kenya | 6 | 2012 | 2018 | 562 (67) | 19.9 (0.3) |
| NZ | Waitati South Island | New Zealand | 2 | 2012 | 2018 | 689 (0) | 10.3 (0) |
| SA | Loskopdam | South Africa | 6 | 2012 | 2017 | 699 (96) | 16.6 (1) |
| USA | Fort Keogh | USA | 6 | 2016 | 2017 | 333 (16) | 8.9 (0.1) |

**Table S2** Three groups of variables used as predictors of bacterial and fungal community composition.

| **1. Broad-scale variables** -  *vary strongly at the large scale* | **2. Ecosystem fertility-related variables** - *vary at the large and the regional scale* | **3. Community-related variable -** *varies at the large and the regional scale* |
| --- | --- | --- |
| Mean annual precipitation (MAP) | Soil organic matter (SOM) | Plant community composition |
| Mean annual temperature (MAT) | Cation exchange capacity (CEC) |  |
| Nitrogen (N) deposition | pH |  |
| Geographical distances | Base saturation (BS) |  |
|  | Carbon to nitrogen ratio (C:N) |  |
|  | Total nitrogen (N) |  |
|  | Total phosphorus (P) |  |
|  | Available phosphorus (P Olsen) |  |
|  | Sand |  |
|  | Silt |  |
|  | Clay |  |
|  | Extractable calcium (Ca) |  |
|  | Extractable magnesium (Mg) |  |
|  | Extractable potassium (K) |  |
|  | Plant biomass |  |

**Table S3** Large-scale predictors of bacterial and fungal community distances. The table is broken into three variable groups. The upper part of the table shows the broad-scale variables and the ecosystem fertility-related variables that had a significant effect in the MRM model. R^2^ values of the individual relationships between the variables and bacterial/fungal community composition are shown together with the coefficients and P values for each of the variables in the MRM model. The coefficients of variables shaded in grey were used to weigh these variables before summing them to obtain a single representative composite environmental variable in Fig. 3. The third group (bottom row) shows the variation in microbial communities explained by plant community composition alone.

|  |  | **Bacterial community composition** | | | **Fungal community composition** | | |
| --- | --- | --- | --- | --- | --- | --- | --- |
|  |  | Individual R^2^ | Coefficient (model) | P-value (model) | Individual R^2^ | Coefficient (model) | P-value (model) |
| **1. Broad-scale**  **variables** | Geo. dist. | 0.04 | 0.020 | 0.002 | 0.15 | 0.003 | 0.001 |
|  | MAT | 0.03 | 0.010 | 0.003 | 0.13 | 0.020 | 0.001 |
|  | MAP | - | - | - | 0.03 | 0.006 | 0.001 |
|  | N deposition | 0.18 | 0.040 | 0.001 | 0.13 | 0.020 | 0.001 |
| **2. Ecosystem fertility-related variables** | Plant biomass | 0.02 | 0.010 | 0.001 | 0.05 | 0.006 | 0.008 |
|  | pH | 0.27 | 0.040 | 0.001 | 0.09 | 0.020 | 0.001 |
|  | CEC | 0.16 | 0.020 | 0.001 | 0.06 | 0.010 | 0.001 |
|  | N total | 0.10 | 0.020 | 0.001 | - | - | - |
|  | C:N | - | - | - | 0.02 | 0.005 | 0.013 |
|  | BS | 0.34 | 0.040 | 0.001 | - | - | - |
|  | Sand | 0.05 | 0.010 | 0.001 | 0.02 | 0.006 | 0.008 |
| **1 & 2** | **Full model R^2^** | **0.65** |  |  | **0.44** |  |  |
| **3. Community-**  **related variable** | Plant community composition | **0.27** |  |  | **0.51** |  |  |

**Table S4** The percentage of reads belonging to different bacterial and fungal phyla in the study

| Bacterial phyla | % of total reads | Fungal phyla | % of total reads |
| --- | --- | --- | --- |
| Actinobacteria | 41.6730 | Ascomycota | 62.3605 |
| Firmicutes | 16.4703 | Basidiomycota | 20.6830 |
| Proteobacteria | 13.7921 | Glomeromycota | 7.2779 |
| Chloroflexi | 7.8897 | Unknown | 5.8253 |
| Acidobacteria | 6.2957 | Mortierellomycota | 2.4193 |
| Verrucomicrobia | 3.0793 | Chytridiomycota | 0.4171 |
| Bacteroidetes | 2.9213 | Olpidiomycota | 0.4162 |
| Planctomycetes | 2.2374 | Rozellomycota | 0.2707 |
| Thaumarchaeota | 1.6295 | Entorrhizomycota | 0.1831 |
| Gemmatimonadetes | 1.5125 | Mucoromycota | 0.0627 |
| Tectomicrobia | 0.5941 | Zoopagomycota | 0.0324 |
| Saccharibacteria | 0.5819 | Blastocladiomycota | 0.0140 |
| Armatimonadetes | 0.4865 | Kickxellomycota | 0.0093 |
| Nitrospirae | 0.1723 | Aphelidiomycota | 0.0077 |
| Elusimicrobia | 0.1121 | Monoblepharomycota | 0.0071 |
| Cyanobacteria | 0.0676 | Entomophthoromycota | 0.0067 |
| Latescibacteria | 0.0610 | Basidiobolomycota | 0.0038 |
| Unknown | 0.0586 | Fungi_phy_Incertae_sedis | 0.0021 |
| Parcubacteria | 0.0549 | Calcarisporiellomycota | 0.0011 |
| FBP | 0.0522 |  |  |
| Chlorobi | 0.0393 |  |  |
| Ignavibacteriae | 0.0362 |  |  |
| TM6 | 0.0292 |  |  |
| Fibrobacteres | 0.0256 |  |  |
| Euryarchaeota* | 0.0207 |  |  |
| Omnitrophica | 0.0165 |  |  |
| Chlamydiae | 0.0139 |  |  |
| Candidatus | 0.0118 |  |  |
| BRC1 | 0.0101 |  |  |
| Deinococcus.Thermus | 0.0094 |  |  |
| Spirochaetae | 0.0083 |  |  |
| FCPU426 | 0.0083 |  |  |
| Peregrinibacteria | 0.0058 |  |  |
| Woesearchaeota* | 0.0044 |  |  |
| Microgenomates | 0.0041 |  |  |
| BJ.169 | 0.0027 |  |  |
| WS2 | 0.0023 |  |  |
| Tenericutes | 0.0015 |  |  |
| Hydrogenedentes | 0.0011 |  |  |
| GAL15 | 0.0010 |  |  |
| RBG.1 | 0.0009 |  |  |
| WWE3 | 0.0008 |  |  |
| Fusobacteria | 0.0007 |  |  |
| CPR2 | 0.0005 |  |  |
| Gracilibacteria | 0.0004 |  |  |
| SBR1093 | 0.0004 |  |  |
| Bathyarchaeota* | 0.0004 |  |  |
| Nitrospinae | 0.0003 |  |  |
| Aenigmarchaeota* | 0.0003 |  |  |
| SR1 | 0.0003 |  |  |
| Deferribacteres | 0.0002 |  |  |
| Parvarchaeota* | 0.0002 |  |  |
| Lentisphaerae | 0.0001 |  |  |
| PAUC34f | 0.0001 |  |  |
| Aminicenantes | 0.0000 |  |  |
| WS1 | 0.0000 |  |  |
| Synergistetes | 0.0000 |  |  |

* Archaea

**Appendix S1** Extended materials and methods

*Analyses of soil properties*

SOM [%] was calculated as the loss of dry matter at 550°C expressed as a percentage of dry matter (Heiri *et al.* 2001). Total soil N and total C [%] were determined on ground soil, dried 48h at 70°C, on a Flash 2000 CN analyser (ThermoFisher Scientific, Waltham, MA, USA). Total P [ppm] was determined using acid digestion with H_2_SO_4_, salicylic acid, H_2_O_2_ and selenium (Novozamsky *et al.* 1983). Available P [ppm] was analysed following the Olsen extraction method (Olsen S, Cole C, Watanabe F 1954) using a Continuous Flow Analyser (CFA) SAN++ (Skalar, Breda, The Netherlands). CEC [meq/100g] and BS [%] were estimated based on the exchangeable H^+^ and total exchangeable bases – TEB [meq/100g]. For this, cations in ammonium acetate extract were measured following Reeuwijk (2002) using an Inductively Coupled Plasma Spectrophotometer (ThermoFisher Scientific, Waltham, MA, USA) and acidity was determined following Brown (1943). pH of air-dried soil was measured using a pH meter (Hanna HI 3222; Hanna Instruments, Woonsocket, RI, USA) in 1:2.5 w:v soil in 1M KCl suspension (Blakemore *et al.* 1987). Soil texture was analysed by determining the percentage of primary particles (sand: 2000-53 µm, silt: 53-2.0 µm, and clay: < 2.0 µm), following the method of Gee & Bauder (1986).

*PCR conditions and library preparation*

Each 25 µl reaction mixture contained 2 µl of the sample, 0.5 µM of each forward and reverse primer, 1X PCR buffer, 200 µM dNTPs and 1 U Phusion High-Fidelity DNA polymerase (New England Biolabs, Ipswich, MA, USA). PCR conditions were as follows: initial denaturation at 98 °C for 60 s, followed by 30 (35 for fungi) cycles of: denaturation at 98 °C for 30 s, annealing at 55 °C for 30 s, extension at 72 °C for 30 s; and an additional extension of 72 °C for 10 min. The success of amplification was tested on 1.5% agarose gel. For the samples that did not amplify successfully, amplification was attempted again with a modified mixture that contained 2 µl of the sample and 1 µM of forward and reverse primer. Successful PCR products were diluted 50-fold and a second PCR was performed using dual barcoded primers with Illumina adapters (2.5 µl of diluted PCR products and 0.1 µM of each primer). The conditions were: 98 °C for 60 s, 12 cycles: at 98 °C for 10 s, 63 °C for 30 s, 72 °C for 30 s; and 72 °C for 5 min. PCR products were run on an agarose gel and successful amplicons were purified and normalized using the SequalPrep Normalization Plate Kit (ThermoFisher Scientific) and pooled into a single library. The library was purified using QIAquick Gel Extraction Kit (Qiagen, Venlo, the Netherlands) and quantified using qPCR (KAPA Library Quantification Kits, Kapa Biosystems, Wilmington, MA, USA).

*Bioinformatics analyses, quality filtering and functional annotation of fungal sequences*

After trimming to 280 bp and 250 bp for bacteria and fungi respectively, the paired-end reads were merged and primers were removed. This trim length was chosen because it was the optimal length for merging paired reads by removing reduced-quality bases at the end. Merged sequences were quality filtered using the expected number of errors (E) as a measure of read quality, with a threshold of E_max_ = 0.5. This yielded 10.8 M and 4.02 M of good-quality reads, for bacteria and fungi, respectively. Following singleton removal, the sequences were clustered into OTUs based on 97% similarity using the UPARSE-OTU algorithm (Edgar 2013) which automatically detects and filters out chimaeras. Filtered reads were then mapped to the OTUs with an identity threshold of 0.97, yielding an OTU table for bacteria and fungi.

The number of reads per subsample was rarefied to 6,046 for bacteria and 1,231 reads for fungi. Most bacterial subsamples had more sequences than the chosen rarefaction depth but 60 samples had fewer sequences than this threshold. Out of these, 13 samples had too few sequences or were clear outliers and they were therefore discarded and 47 were upsampled to contain 6,046 sequences while retaining the original proportions (as in De Gruyter et al. (2020)), leaving 402 bacterial samples. Given that we were interested in plot-level community composition, this procedure was done in cases when it could be verified that the upsampled bacterial communities do not notably deviate from those in other subsamples of their group (i.e. those sampled from the same plot) which demonstrated that their overall quality was not compromised. For fungi, 13 samples had fewer sequences than the chosen rarefaction depth, and they were omitted leaving 402 samples in total. Both for bacteria and fungi, at least 3 samples per plot were retained.

To annotate *sintax*-assigned fungal sequences to known genera in the UNITE database, we used NCBI’s BLAST algorithm with default settings. OTUs were then assigned to particular taxa if they had a maximum E-value of 10^-36^ and from this, the lowest E-value hit with a known genus was selected. If there were none, the genus level was left unassigned. OTUs were subsequently assigned to functional groups if the genus was successfully matched with one of the genera with known lifestyles in Tedersoo et al. (2014) and Liu et al. (2016). The functional groups included: saprotrophs, plant pathogens, animal parasites, mycoparasites, arbuscular mycorrhizal fungi and ectomycorrhizal fungi.

*Microbial abundance analyses using qPCR*

Each 20 µl reaction mixture contained 4 µl of the sample, 0.5 µM of each forward and reverse primer, 1 x ROX high and 10 µl of KAPA SYBR FAST qPCR master mix (Kapa Biosystems, Wilmington, MA, USA). qPCR conditions were as follows: initial denaturation at 95°C for 3 min, followed by 40 cycles of: denaturation at 95 °C for 3 s, annealing at 57 °C (52 °C for fungi) for 20 s, extension at 72 °C for 12 s; finishing with 35 s at 50 °C. Prior gel-electrophoresis with these primers and reaction conditions showed the reactions were highly specific. Melting curve analysis of all amplicons was conducted to confirm that fluorescence signals originated from specific amplicons and not from primer-dimers or other artefacts. Standard curves were generated using duplicates of 10-fold dilutions of amplicons derived using the same primers, isolated from gel using QIAquick Gel Extraction Kit (Qiagen, Venlo, the Netherlands) and quantified using Qubit fluorometer (Invitrogen GmbH, Karlsruhe, Germany). Fungal and bacterial gene copy numbers were derived from a regression equation based on the standard curves (with minimal R^2^ > 0.99) by relating the quantification cycle (Cq) value of each sample to the Cq values of standards with the known number of copies. All reactions were performed in duplicate and the number of bacterial and fungal copies was then averaged (a deviation of Cq between replicates < 1 was used as a passing criterion) and expressed per g of soil dry weight.

**Appendix S2** – Microbial community composition predictors at different productivity levels

At high productivity, broad-scale variables and ecosystem fertility-related variables explained 72% of the variation in bacterial community composition, and the most influential individual predictor was base saturation explaining 35% of the variation. Plant communities explained 34% of the variation alone and when added to the other variables the amount of the variation explained increased from 72% to 77%. Broad-scale variables and ecosystem fertility-related variables explained 82% of variation at low productivity, where the most important individual factor was again base saturation (R^2^ = 0.58). Plant community composition added 5% of variation when included in the model with other variables (and alone it explained 34% of the variation).

At high productivity, broad-scale variables and ecosystem fertility-related variables explained 55% of the variation in fungal community composition and the most influential individual predictor was geographical distance (R^2^ = 0.24). Plant communities alone explained 67% of the variation and when added to the above model, the total variance explained increased from 55% to 76%. At low productivity, 48% of the variation was explained by the model with broad-scale variables and ecosystem fertility-related variables, where geographical distance explained the most individual variation (R^2^ = 0.18). Plant community composition alone explained 78% of the variation and when added to the model with broad-scale variables and the ecosystem fertility-related variables, the total amount of variation increased from 48% to 78%.
